# A model of Zebrafish Avatar for co-clinical trials

**DOI:** 10.1101/784041

**Authors:** Alice Usai, Gregorio Di Franco, Patrizia Colucci, Luca Emanuele Pollina, Enrico Vasile, Niccola Funel, Matteo Palmeri, Luciana Dente, Alfredo Falcone, Luca Morelli, Vittoria Raffa

**Affiliations:** Department of Biology, Università di Pisa, S.S. 12 Abetone e Brennero 4, 56127 Pisa, Italy.; Department of Traslational Research and of New Surgical and Medical Technologies, General Surgery Unit, University of Pisa, Via Paradisa 2, 56124 Pisa, Italy.; Department of Surgical, Medical, Molecular Pathology and Critical Area, Division of Surgical Pathology, University of Pisa,Via Paradisa 2, 56124 Pisa, Italy.; Division of Medical Oncology, Pisa University Hospital, Via Roma 67, 56126 Pisa, Italy.

**Keywords:** patient-derived xenografts, zebrafish, chemosensitivity, equivalent dose, translational research

## Abstract

Animal ‘‘Avatars’’ and co-clinical trials represent an emerging concept for implementing schemes of personalized medicine in oncology. In a co-clinical trial, the cancer cells of the patient tumor are xenotransplanted in the animal Avatar for drug efficacy studies and data collected in the animal trial are used to plan the best drug treatment in the patient trial. Recently, zebrafish has been proposed for implementing Avatar models but the lack of a general criterion for chemotherapy dose conversion from humans to fishes represents a limitation for conducting co-clinical trials.

Here, we validate a simple, reliant and cost-effective Avatar model based on the use of zebrafish larvae; by crossing data from safety and efficacy studies, we found a basic formula for the estimation of the dose to be used for running co-clinical trials and we validate it in a clinical study enrolling 24 adult patients with solid cancers (XenoZ, NCT03668418).

## INTRODUCTION

Precision medicine refers to the approaches for tailoring a medical treatment to the individual characteristics of each patient (1). In particular, the “Mouse Avatar” is an emerging approach of precision medicine in oncology that has recently grown in importance (2); it implicates the xenotransplantation of cancer cells from patient tumor sample in mouse models to use them in drug efficacy studies. Mouse Avatars can be used to run “co-clinical trials” (3). In a co-clinical trial, the patient and murine trials are concurrently conducted and the drug efficacy response of the mouse study provides data to plan the best drug treatment of the patient tumor (4). The advantage of this approach is that each patient has his/her own tumor growing in an *in vivo* system, thereby allowing the identification of a personalized therapeutic approach. Nowadays, there are companies providing mouse Avatar generation and drug testing services to patients at a cost of tens thousands of dollars (5). The high cost is directly associated to the time-consuming process and the requirement of immunosuppressed strains (6). Unfortunately, this makes Avatars a cutting-edge technology available only for few people, posing a serious threat to the equal right to health for everyone. Recently, it has been proposed the use of zebrafish to make Avatars available for every patient and the approach sustainable for National Healthcare Systems. Zebrafish cancer models overcome the drawbacks of xenografts in mice (7). Zebrafish is highly fecund, develops rapidly and requires simple and inexpensive housing. Zebrafish embryos are transparent, allowing to image the engrafted cells *in vivo*, and they have high permeability to small molecules such as drugs used for chemotherapy. Last but not least they have low ethical impact when used in the larval stage from fecundation to 120 hours post fertilization (hpf) (8). Zebrafish larvae as model for human cancer cell xenografts have been firstly reported in 2005 (9). Since then, the use of zebrafish *in vivo* model of xenotransplantation has increased considerably (10). Several human cancer cells lines e.g. melanoma, glioma, adenocarcinoma, breast, pancreas and prostate cancer cell lines (11) as well as fragments of human cancer tissues (12) have been tested to date in zebrafish as engraftment host. Larvae provide a rejection-free permissive environment, where the xenotransplanted human cancer cells rapidly proliferate, migrate, form masses and induce neo-angiogenesis, after injection (13). Most importantly, zebrafish larvae xenografts provide similar chemosensitive response of mouse xenografts (14).

However, in order to move forward in new paradigm of co-clinical trial using zebrafish Avatars, some critical aspects need to be solved. The biggest issue is related to the lack of the “equivalent dose” for translating the chemotherapy dosage used in humans to zebrafish larvae because one cannot apply the interspecies allometric approach for dose conversion from human to animal. The caveat is that chemotherapy drugs have to be administered in the fish water rather than injected as parenteral formulations. Therefore, drug safety and efficacy assessments are necessary to estimate the equivalent dose to administer (15). The present study aims to fill the gap regarding the dose conversion between zebrafish larvae and humans. A safety/efficacy study has been carried out in HCT 116 and MIA PaCa-2 cancer cell lines by testing 10 different chemotherapy regimens used in cancer treatment, i.e. FOLFOX (5-Fluorouracil + Lederfolin + Oxaliplatin), FOLFIRI (5-Fluorouracil + Lederfolin + Irinotecan), FOLFOXIRI (5-Fluorouracil + Lederfolin + Oxaliplatin + Irinotecan), ECF (5-Fluorouracil + Cisplatin + Epirubicin), FLOT (5-Fluorouracil + Lederfolin + Oxaliplatin + Docetaxel), GEMCIS (Gemcitabine + Cisplatin), Gem/nab-P (Gemcitabine + nab-Paclitaxel), GEMOX (Gemcitabine + Oxaliplatin), Gemcitabine, 5-Fluorouracil. We found a general criterion for dose equivalence that has been validated on zebrafish Avatar receiving fresh tissue fragments taken from surgical specimens of patients underwent surgical operation for hepato-biliary-pancreatic cancer and gastro-intestinal cancer.

## RESULTS

### Zebrafish safety study

Dose-response analysis for the determination of the effects of chemotherapy treatment on larvae was based on the evaluation of the phenotype resulting from the exposure (i.e. normal, aberrant and dead). In particular, we exposed larvae to 10 different chemotherapy treatments (GEM, GEMOX, GEM/nab-P, GEMCIS, 5-FU, FOLFOX, FOLFIRI, FLOT, FOLFOXIRI, ECF, see supplementary tables S1 and S2) for 72 hours, from 48 to 120 hpf (Figure 1). Chemotherapy treatments induced death and a variety of malformations in larvae, including yolk sac edema, pericardial edema and spine deformation. For all regimens, deviation from phenotype without defect (normal phenotype) increased with the increase of drug concentration. Linear regression analysis showed an excellent relationship between the linear or logarithmic concentration of the chemotherapy drug and the incidence of normal phenotype (R^2^>0.95; p<0.05 for any protocol tested) or the incidence of mortality (R^2^>0.87; p<0.05 for any protocol tested), (Figure 1). For any chemotherapy treatment, the dose that is lethal to 25% of the population (LD25) and the concentration at which 50% of the normal phenotype is inhibited (IC50) was determined (Figure 2A). Such data were also expressed as Conversion Factor (CF):

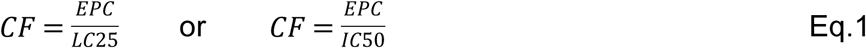

With EPC defined as human Equivalent Plasma Concentration, given by:

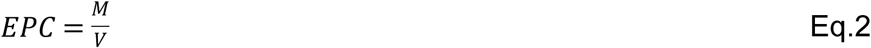

*M* being the total amount (mg) of chemotherapy administered to humans by the clinicians involved in the present study, *V* (ml) being the mean volume of human plasma (the EPC value for each regimen is given in table S2).

**Figure 1.**
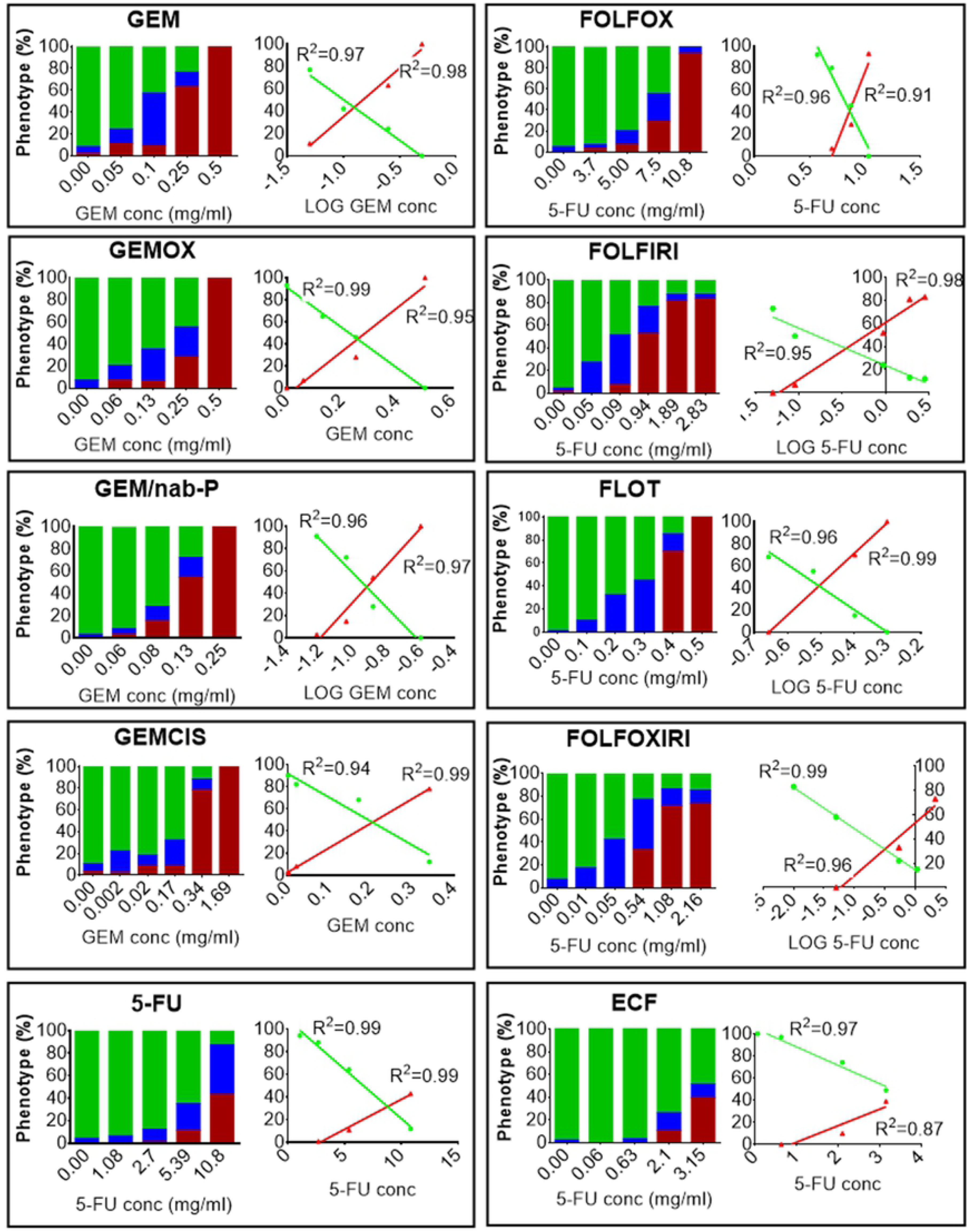
Chemotherapy toxicity study. Zebrafish embryos 2 dpf were incubated with media (E3 supplemented with 10000 U/ml penicillin and 100 μg/ml streptomycin) modified with chemotherapy drugs or not modified at 35°C for 3 days. At the end of the treatment the percentage of dead embryo (red), aberrant (blue) and normal phenotype (green) was evaluated after fixation and stereomicroscope observation. In GEM, GEMOX, GEM/nab-P, GEMCIS we reported the Gemcitabine concentration in the x-axis. In 5-FU, FOLFOX, FOLFIRI, FLOT FOLFOXIRI, ECF we reported the 5-Fluorouracil concentration in the x-axis. Control group showed an alteration from normal phenotype ≤10%. For each chemotherapy regimen a dose-response and the relative linear regression analysis of the normal phenotype and the dead embryos are shown. The resulting R square is reported. The results presented are a pool from three independent biological replicates (n=90).

**Figure 2.**
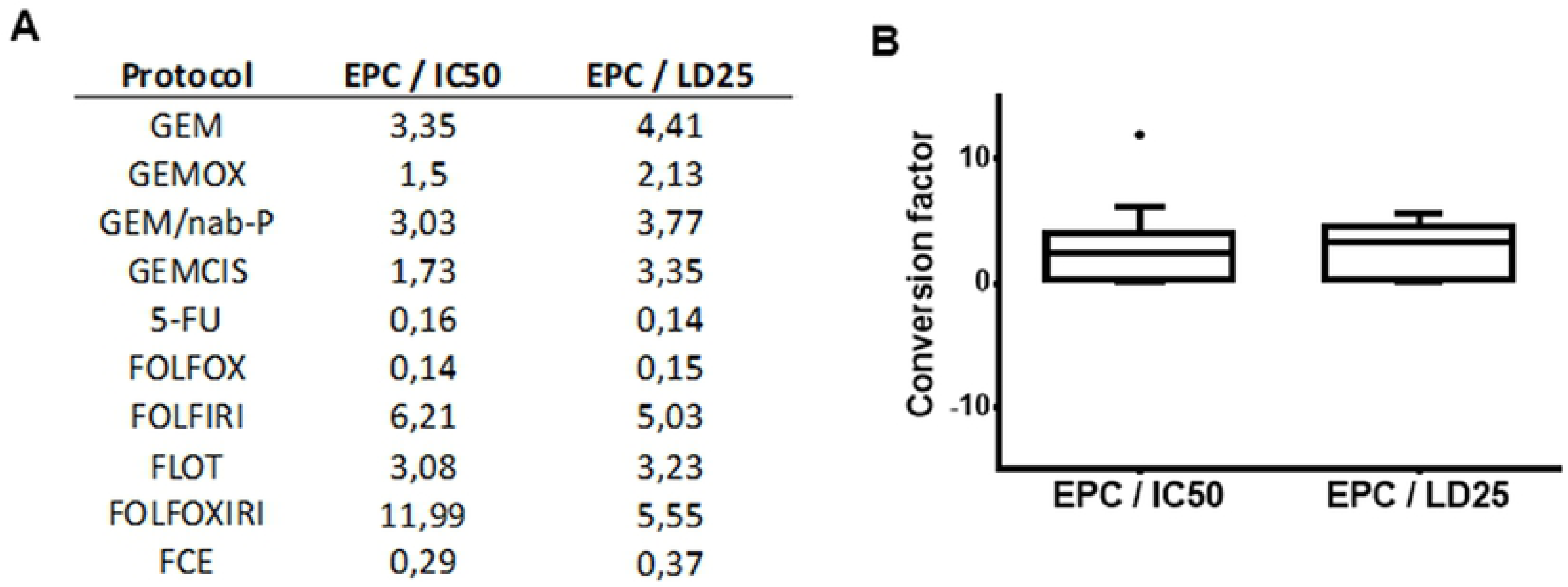
Estimation of the maximum tolerated dose. (A) Table and (B) box plot displaying EPC/IC50 EPC/LD25 ratios for all chemotherapy protocols.

In the present study, we fixed the 75 percentiles of the box plots (Figure 2B) as CF corresponding to the maximum tolerated doses (MTDs). Interestingly, this value was similar for the 2 conditions, i.e. CF=4.1 and CF=4.6 for EPC/LD25 and EPC/IC50, respectively. Consequently, the toxicity study established a conversion factor 4.6 ≤ CF < ∞ as the range for the determination of the equivalent dose required for running co-clinical trials with an acceptable safety level for the zebrafish trial.

### Zebrafish efficacy study

For the estimation of the equivalent dose, we conducted an *in vivo* efficacy study based on human cancer cell lines, whose chemosensitivity has been already characterized in the literature. Specifically, 2 dpf larvae were xenotransplanted with DiI-stained human colorectal carcinoma cell line (HCT 116) or human pancreatic carcinoma cell line (MIA PaCa-2) into the yolk sac. To confirm the presence of the xenograft, injected larvae were screened by fluorescence microscopy 2 hours post injection (hpi). The screened larvae were randomly distributed in a multiwell plate (1 embryo/well) and equally divided among groups (control and chemotherapy regimens). In absence of chemotherapy, the DiI-stained area shows a statistically significant increase over the time (Figure 3, control group) and the block or the inversion of this tendency has been considered in the present study as a hallmark of chemotherapy effect. Indeed, we tested 4 chemotherapy regimens (5-FU, FOLFOX, FOLFOXIRI, FOLFIRI), which are the standard of care for the treatment of colorectal cancers, on HCT 116 cells xenotransplanted in 2 dpf larvae. According to the toxicity study, we used conversion factors CF>4.6. First, a CF=8 was tested but data showed a statistically significant increase of the DiI-stained area at 1 dpi and 2 dpi for all the regimens, suggesting the inefficacy of chemotherapy treatment at the CF used (Figure 3A). Therefore, we tested chemotherapy protocols at a higher concentration, corresponding to CF=5. Interestingly, FOLFOXIRI were found to inhibit the increase of the stained area at 1 dpi and 2 dpi (p>0.05), as opposite to the control, 5-FU, FOLFOX and FOLFIRI that showed a statistically significant progression (Figure 3B). The effect of FOLFOXIRI treatment was also confirmed by quantification of apoptosis in xenotransplanted (DiI-positive) cells revealing a significant increase of pyknotic nuclei with respect to the control group (no chemotherapy drugs) (Figure S1A).

**Figure 3.**
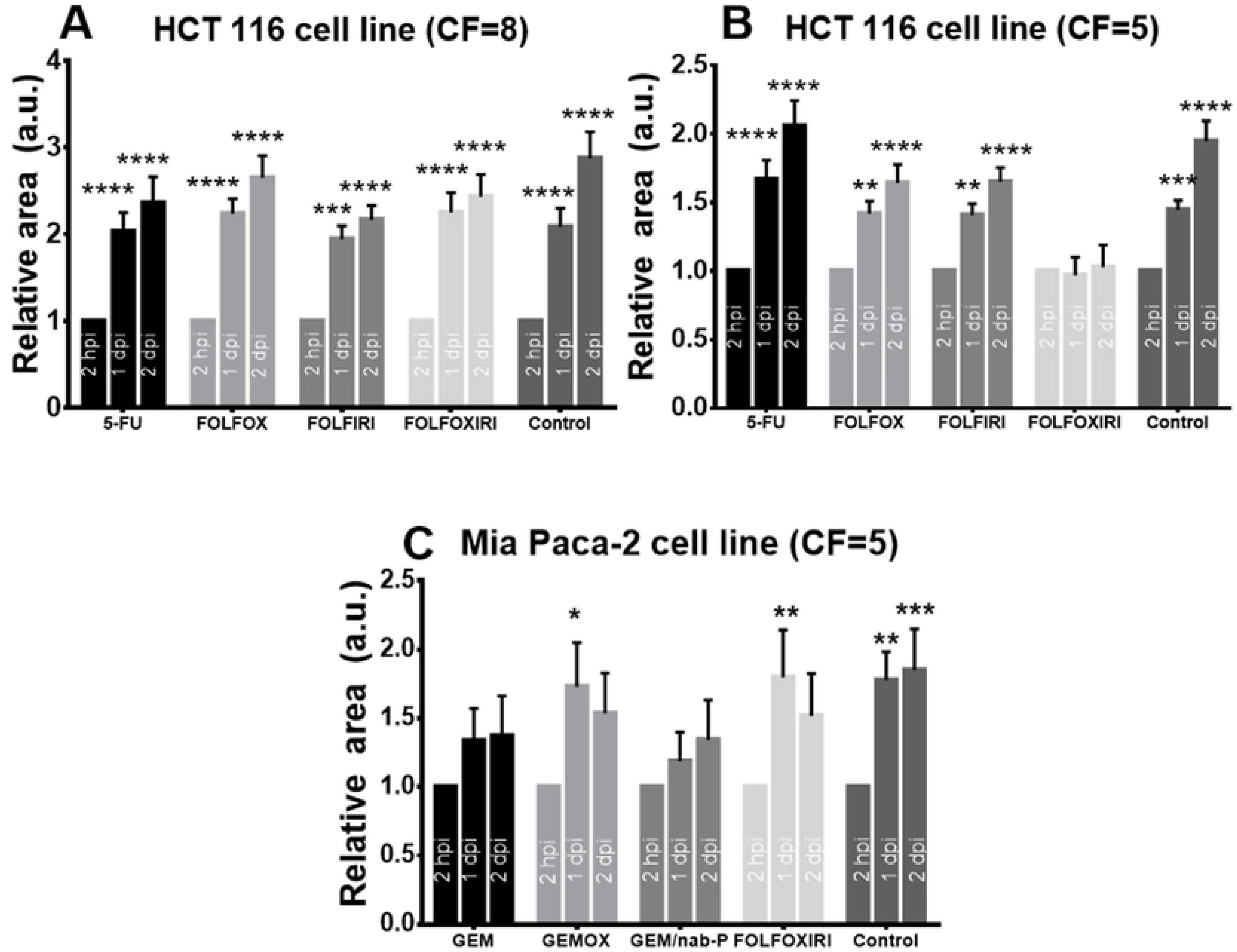
Efficacy analysis. Evaluation of the effects of chemotherapy on cancer cell lines (HCT 116, MIA PaCa-2) xenotransplanted in 2dpf zebrafish embryos. Each embryo was imaged at 2 hpi, 1 dpi, 2 dpi and the relative area is the Dil-stained area normalized with respect to the 2 hpi time point. (A) Chemosensitivity of HCT 116 xenografts, CF=8. A statistically significant increase of relative area was observed in all groups. (B) Chemosensitivity of HCT 116 xenografts, CF=5. A statistically significant increase of relative area was observed in control, 5-FU, FOLFOX and FOLFIRI but not in FOLFOXIRI. (C) Chemosensitivity of MIA PaCa-2 xenografts, CF=5. A statistically significant increase of relative area was observed in control, GEMOX and FOLFOXIRI treatments but not in GEM and GEM/nab-P. Data are mean ± SEM and representative of three independent assays. n≥15 (embryos), 2-way ANOVA followed by Bonferroni correction (all groups compared against k group). * p < 0.05; ** p < 0.01; *** p< 0.001; **** p< 0.0001.

The next step was to confirm the value of CF=5 for dose equivalence by testing its efficacy on a different model, i.e. xenotransplanted larvae receiving MIA PaCa-2 cell line. Indeed, we tested 4 chemotherapy regimens (GEM, GEMOX, GEM-Nab, FOLFOXIRI), which are the standard of care for the treatment of pancreatic cancers, on MIA PaCa-2 cells xenotransplanted in 2 dpf larvae. GEM and GEM-Nab-P proved to be the most efficient regimens, with no statistically significant increase of the Dil-stained area at 1 dpi and 2 dpi, as opposite to the control, GEMOX and FOLFOXIRI (Figure 3C).

### Zebrafish Avatar

A total of 6 patients operated for adenocarcinoma of the colon (n=3), pancreatic ductal adenocarcinoma (n=1), and gastric adenocarcinoma (n=2) have been enrolled in the study (NCT03668418) to establish the zebrafish Avatar model. In order to preserve the tumor micro-environment, we decided to xenotransplant fresh tissue fragments screened by the histopathology unity of the Azienda Ospedaliera Universitaria Pisana, by modifying the protocol published by Marques *et al.* (12). Briefly, the tissue was DiI/DiO-stained, disaggregated using Dumont forceps (No.5) into a relative size of ½-¼ the size of the yolk and xenotransplanted in the yolk of 2 dpf larvae. After transplantation, larvae were incubated for 2 h at 35°C, then screened to check for presence of the stained tissue and imaged at 2 hpi, 1 dpi and 2 dpi. Cell engraftment was also confirmed by histological analysis performed at 2 dpi. Hoechst staining documents healthy cell nuclei (Figure S1B) and H&E staining shows the presence of cancer cells that have the typical round-shape morphology with large nuclei (Figure S2A). Interestingly, H&E staining performed on zebrafish embryos xenotransplanted with fragments of normal tissue taken from normal mucosa or pancreatic parenchyma of the surgical specimen do not show any cell with typical cancer morphology but exclusively cells with a typical fibroblast-like shape (Figure S2B). In order to perform the analysis, we measured the size of the region of interest (ROI) corresponding to the stained area at 2 hpi, 1 dpi and 2 dpi (Figure 4A3-C3). The mean size of the tumor mass area measured in each time point was normalized with respect to the 2 hpi time point. We found an increase of the stained area versus time in all cases, which was statistically significant at 2 dpi with respect to the time point 2 hpi for 5 patients of 6 (83%, Figure 5). According to this finding, the measure of the size of the relative stained area at 2 dpi has been fixed as primary measure of the study. Sporadically we also detected cancer cell migration (Figure S3).

**Figure 4.**
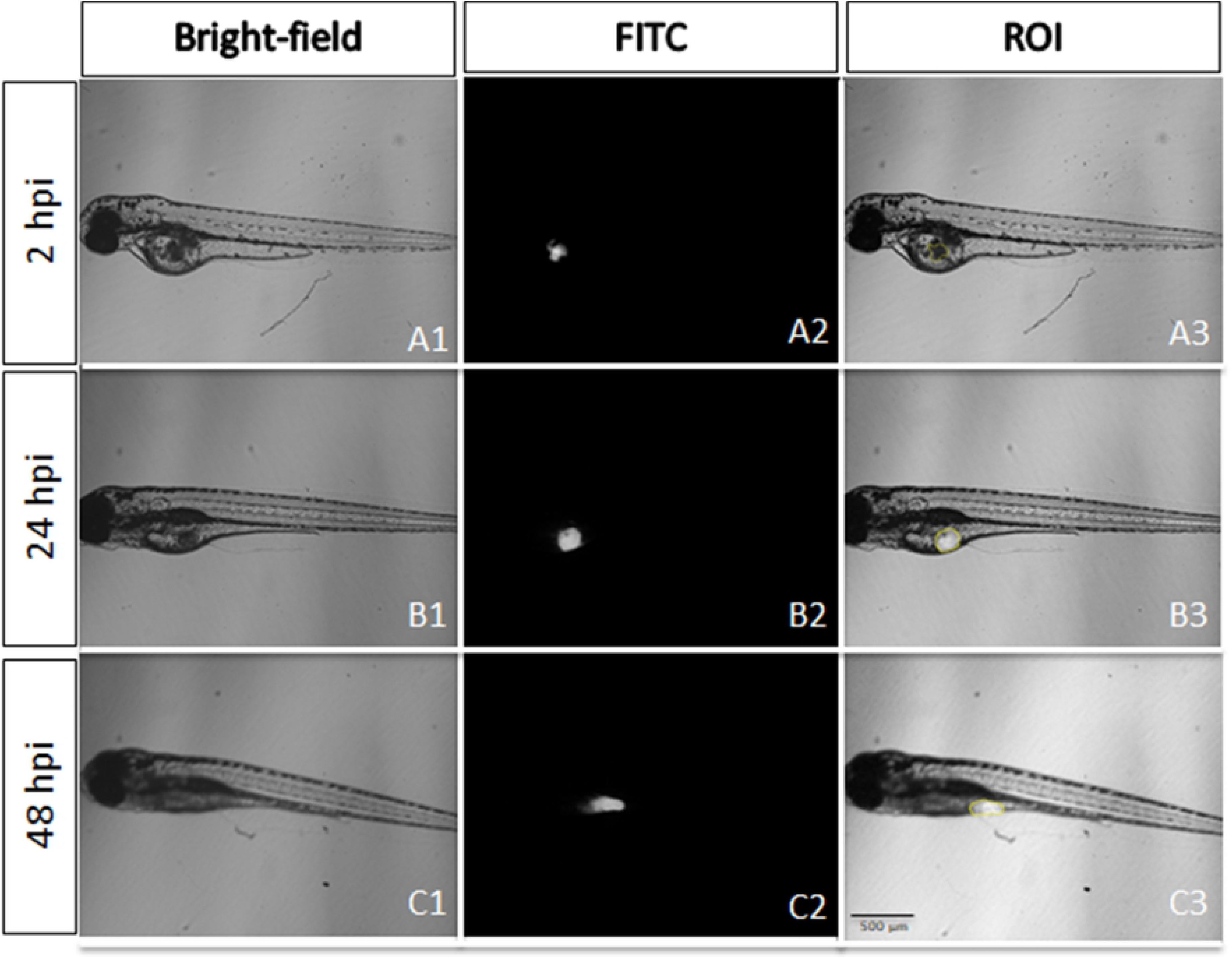
A representative larva xenotransplanted with a fresh tumor specimen of gastric cancer (patient S013). Bright-field images of the grafted larvae (A1-C1), epi-fluorescence images (A2-C2) and overlay (A3-C3), showing the region of interest (ROI; yellow line). All images are oriented so that rostral end is on the left and dorsal end is on the top.

**Figure 5.**
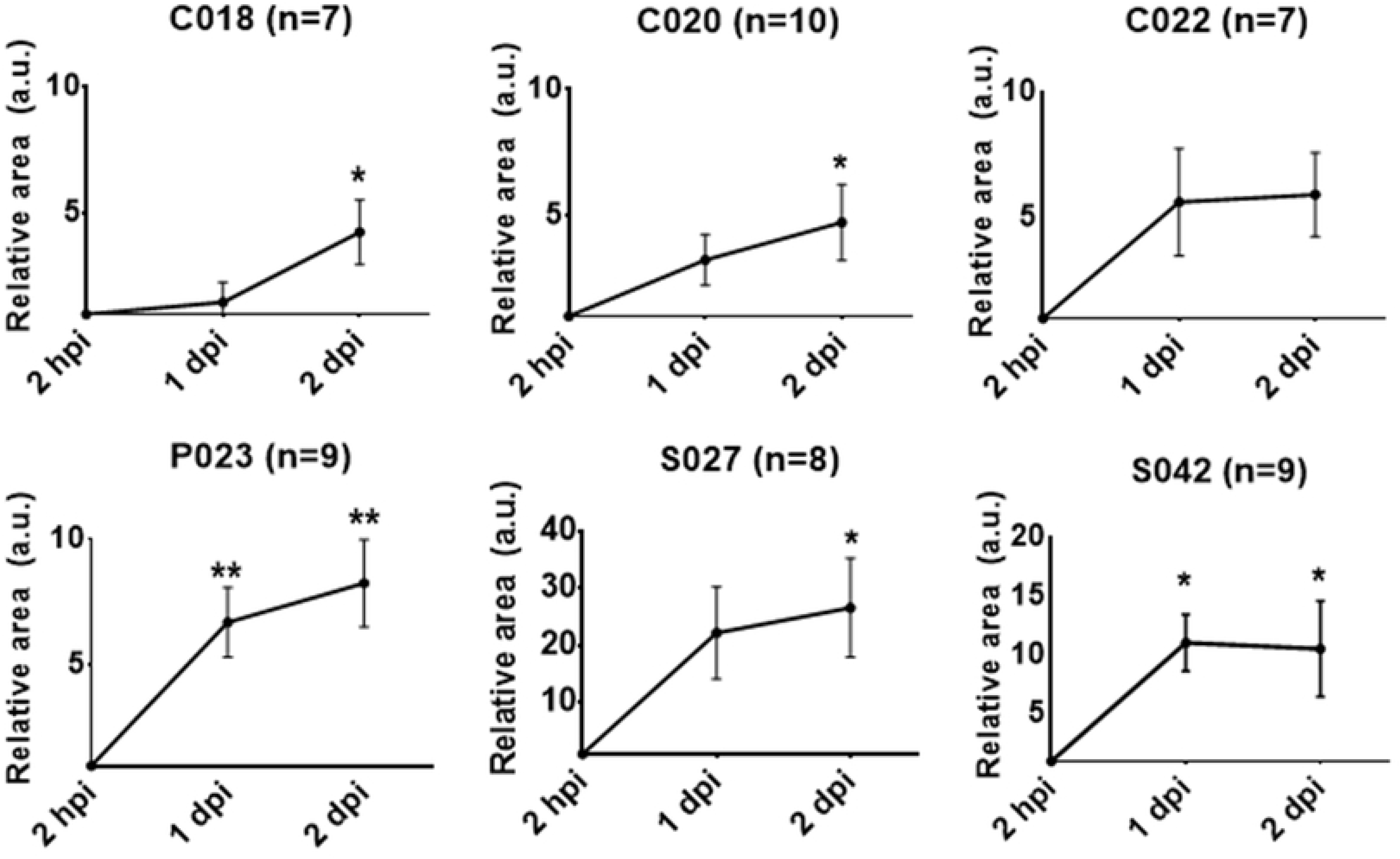
Quantitative analysis of six cases of patient-derived tumor xenografts. Dil-stained area at time point 2 hpi, 1 dpi and 2 dpi was normalized with respect to the time point 2 hpi. Patient enrollment code is reported (C=Colon, P=Pancreas, S=Stomach) and the number of embryos analyzed for each case study is indicated in the image. Results are expressed as mean ± SEM. * p < 0.05; ** p < 0.01 by 1-way ANOVA followed by Bonferroni correction (1 dpi and 2 dpi compared against 2 hpi).

### Zebrafish trial

24 adult patients with pancreatic cancers (n=12), colon cancer (n=8) and gastric cancers (n=4) undergoing a chemotherapy treatment have been recruited for this part of the study. After surgery and histopathology screening, patient biopsies have been xenotransplanted in 100 zebrafish embryos and injected embryos were randomly allocated among 5 groups (4 therapeutic options and one control group). Groups were exposed to all chemotherapy options, according to the cancer type, by dissolving the chemotherapy in fish water, according to the equivalent dose corresponding to CF=5. Two days post treatment the response of zebrafish xenografts to the chemotherapy options was analyzed by monitoring the ROI size at 2 hpi, 1 dpi and 2 dpi (Figure S4). The chemotherapy protocols tested were 5-FU, FOLFOX, FOLFIRI and FOLFOXIRI for colon cancer; GEM, GEMOX, GEM/nab-P and FOLFOXIRI for pancreatic case and FOLFOX, FOLFORI, FLOT and ECF for gastric cancer. We adapted the “Response evaluation criteria in solid tumors (RECIST)” to the fish trial by defining the partial response (PR, at least a 30% decrease in the relative stained area at 2 dpi / 2 hpi, taking as reference the relative stained area at 2 dpi / 2 hpi of the control group) and complete response (CR, at least a 90% decrease in the relative stained area at 2 dpi / 2 hpi, taking as reference the relative stained area at 2 dpi / 2 hpi of the control group) (Figure 6). For patients affected by colon cancer, we observed a PR in 62.5% of patients to FOLFOX, FOLFIRI and FOLFOXIRI but a less frequent response (37.5% of patients) to 5-FU; CR was observed only in a limited number of patients (12.5%) and only to FOLFIRI chemotherapy. For patients affected by pancreatic cancer, we observed a PR to GEM/nab-P (58.33 % of patients), GEM (50%), GEMOX (50%), a limited PR to FOLFOXIRI (33.33 %) but we never observed CR for any chemotherapy treatment. For patients operated for gastric cancer, we observed high incidence of PR to FOLFIRI (100% of patients) but low incidence of PR to FOLFOX, FLOT and ECF (25% of patients); we also observed CR to FOLFIRI in one patient of four.

**Figure 6.**
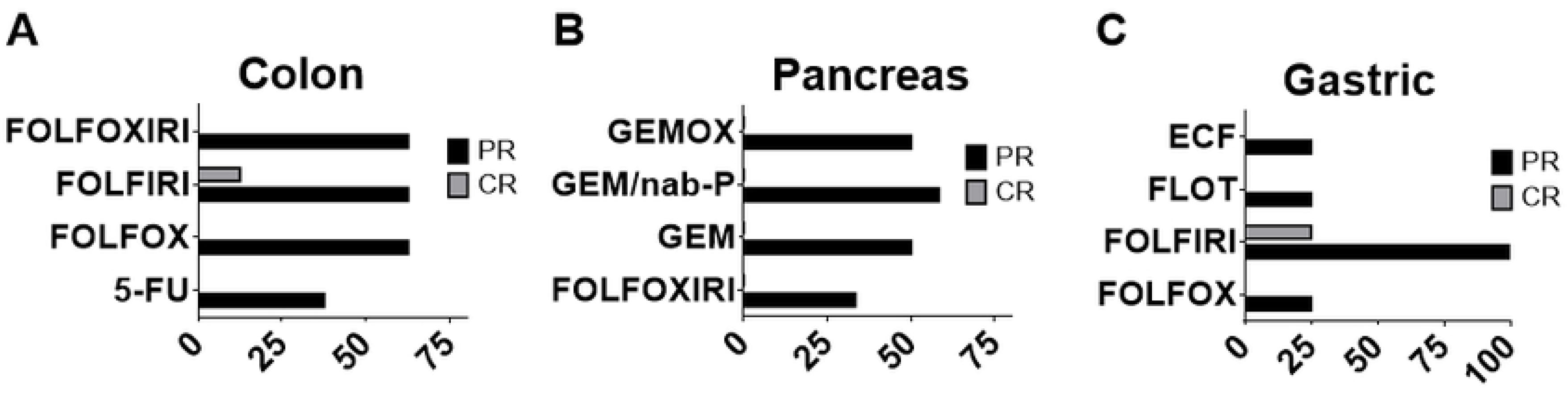
Percentage of Partial Response (PR) and Complete Response (CR). FOLFOXIRI, FOLFIRI, FOLFOX and 5-FU treatments in Zebrafish Avatar xenotransplanted with colon tumor (n=8 patient case analyzed) (A); GEMOX, GEM/nab-P, GEM, FOLFOXIRI treatments in Zebrafish Avatar xenotransplanted with pancreas tumor (n=12 patient case analyzed) (B); ECF, FLOT, FOLFIRI and FOLFOX treatments in Zebrafish Avatar xenotransplanted with gastric tumor (n=4 patient case analyzed) (C).

Interestingly, the zebrafish Avatar can be used to perform chemosensitivity assessment on a single patient basis. Four representative case of patient enrolled in the study are given in Figure 7. As for the two cases of colon cancer, we could observe significant response (p=0.03) to FOLFOXIRI treatment in patient C024, and to 5-FU (p=0.05) and FOLFIRI (p=0.02) in patient C031. As for the two cases of pancreatic cancer, FOLFOXIRI proved to be the efficient regimen in patient P025 (p=0.02) and in patient P030 (p=0.04).

**Figure 7.**
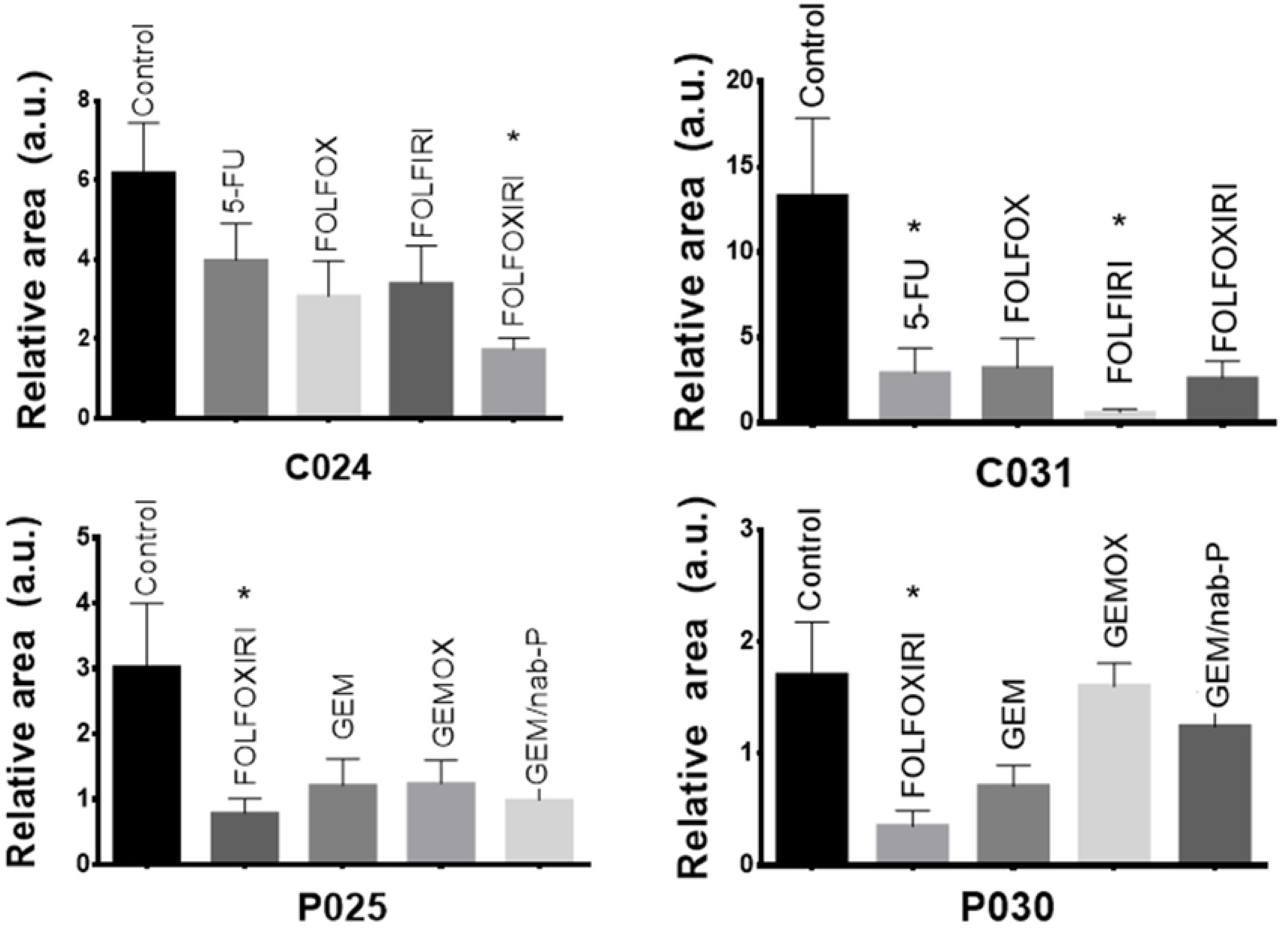
Chemosensitivity assay. 48 hpf embryos were injected with fragments of patient’s tumor tissue and treated for 48 hours with different chemotherapy compounds at the CF=5. Representative cases of colon cancer (patient enrolled code: C024, C031) and pancreatic cancer (patient enrolled code: P025, P030) with quantitative analysis of relative tumor area (2 dpi/2 hpi for colon and 2 dpi/24 hpi for pancreas). All graphs show an increase of the stained area over the time in control group. The cases C024, P025, P030 show a statistically significant regression of the stained area size in FOLFOXIRI treated group. The case C031 shows significant stained area reduction in 5-FU and FOLFIRI treated groups. Results are expressed as mean ± SEM and analyzed by 1-way ANOVA followed by Dunnett’s multiple comparisons test. * p < 0.05, n≥3.

## DISCUSSION

Many studies have demonstrated that preclinical models hold great promise for the implementation of personalized medicine strategies (16–18). Mouse Avatar have been proposed and implemented by different research groups, but still have important practical limitations (2, 19, 20). Zebrafish offers distinct advantages over the murine-based co-clinical trials because of the relatively simple, rapid and cost-effective method to establish a human tumor xenograft model (21). Zebrafish Avatar approach could be used for evaluating individual patient drug responses in a clinically relevant setting or for the high-throughput screening of new molecules. Considering the validity of the Zebrafish Avatar and the affordable costs, the possibility to exploit this model in clinical practice has emerged. To do that, the equivalent dose conversion from human to fish need to be identified.

In this work, we found a general dose conversion criterion based on the following formula:

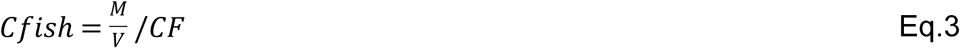

Where *c_fish_*(mg/ml) is the chemotherapy concentration in fish water, *M* is the total amount (mg) of chemotherapy administered to humans, *V* (ml) is the volume of human plasma and *CF* is the conversion factor that we estimated to be *CF*=5. We estimated this value by matching data collected from the safety and efficacy studies performed in zebrafish. The safety study was performed on WT larvae. The efficacy study was performed on larvae xenotransplanted with human cancer cell line whose response to chemotherapy has been already characterized. Specifically, we found that HCT 116 responded to FOLFOXIRI treatment with higher sensitivity, but not to 5-FU, FOLFOX and FOLFIRI at the CF proposed. These results are concordant with the literature suggesting that first-line FOLFOXIRI chemotherapy leads to improved survival and efficacy of metastatic colorectal cancer patient outcomes in comparison to FOLFIRI or FOLFOX chemotherapy (22). We also tested the response of MIA PaCa-2 cells by observing high sensitivity to GEM and GEM/nab-P treatments. This analysis is confirmed by the efficacy data from metastatic pancreatic cancer patients treated with GEM/nab-P (23). Such experimental evidences, obtained by testing the efficacy of chemotherapy on two cancer cell lines from different types of human tumor (colorectal and pancreatic), suggest that the selected therapeutic dose (corresponding to *CF*=5) is effective in killing tumor cells and, in principle, predictive of the best pharmacological treatment. Indeed, we suggest the use of the conversion factor *CF*=5 in any co-clinical trial using zebrafish Avatars. This represents a starting point for any further research step that aims to validate the zebrafish Avatars as valuable tools to support the oncologists in the clinical routine. Potential applications are the evaluation of the disease prognosis and chemosensitivity assays for the prediction of the most effective chemotherapy scheme. Indeed, we validated an approach consisting in the xenotransplantation of pieces of the patient tumor tissue, after surgery and histopathology screening, in order to obtain a model for testing the response of the patient tumor to the different chemotherapy regimens, with an assessment in less than one week (Figure S4). The xenotransplantation of cancer cells isolated and propagated from patient tumors in zebrafish is an approach more popular than the xenotransplantation of tissue fragments. Unfortunately, isolated cancer cells tend to lose cell heterogeneity and the stromal contribution. Moreover, during the process of isolation and adaptation, clones with a higher proliferative rate than that of the primary tumor are selected and thus they could not be representative of the cancer cell population (24). Indeed, for precision medicine and personalized medicine, the xenotransplantation of biopsy or surgical specimen fragments screened by the pathologist would be recommended to develop patient-derived xenografts in which the stromal counterpart and cancer cell heterogeneity are both preserved (25). Our data suggest that fresh tumor tissue transplanted in 2dpf larvae can engraft and survive in the host, as documented by histological analysis showing typical cancer cell morphology (H&E staining, Figure S2A) and absence of pyknotic nuclei (Hoechst staining, Figure S1B). The survival rate of the xenografted host was acceptable, at both 1 dpi (81%, n=101) and at 2 dpi (68%, n=101). We also detected the capacity of cancer cell extravasion and dissemination in distal tissues (Figure S3). As the relative area at 2 dpi/2 hpi has been fixed as primary measure of the study, we performed the efficacy tests under the assumption that a statistically significant decrease of this measure with respect to control group (no chemotherapy) is a hallmark of chemotherapy response. Specifically, we tested chemosensitivity in 24 human tumor fragments taken from surgical specimen. To this purpose, pieces of tumor tissue were microinjected in zebrafish embryos to create Zebrafish Avatar and treated with chemotherapy drugs at a concentration corresponding to *CF*=5. Interestingly, our experimental data have shown good agreement with observations registered in the common clinical practice. In fact, for patients affected by colon cancer (Figure 6A), we found a superiority of the chemotherapy treatment when a combination of drugs are used (FOLFOX, FOLFIRI and FOLFOXIRI) respect to the use of only 5-FU (26). Additionally, we found a similar response to FOLFOX and FOLFIRI (27). The higher aggressiveness of pancreatic cancers associated with a lower response to chemotherapy compared to colon and gastric cancers may be the reason why a complete response was never observed in our experiments for this group of patients (Figure 6B). For the group of patients affected by gastric cancer (Figure 6C), we found an excellent response to FOLFIRI that can be considered an acceptable first-line treatment for advanced gastric cancers (28).

Interestingly, the use of zebrafish Avatars allows to appreciate a different response to different chemotherapy regimens on a single-patient basis (Figure 7). Further tests will be necessary to fully validate the zebrafish Avatar here proposed as a clinical tool predictive of the most effective treatment for each patient. Future experiments will be devoted to enroll in the study a higher number of cases in order to correlate the chemosensitivity results obtained in the animal trial with the response to the chemotherapy treatment observed in the human trial.

## MATERIALS AND METHODS

### Zebrafish husbandry

Zebrafish *(Danio rerio)* were handled in compliance with local animal welfare regulations (authorization n. 99/2012-A, 19.04.2012; authorization for zebrafish breeding for scientific purposes released by the “Comune di Pisa” DN-16/43, 19/01/2015) and standard protocols approved by Italian Ministry of Public Health, in conformity with the Directive 2010/63/EU. Zebrafish fertilized eggs were obtained by natural mating of *wild-type* fishes at our facilities and the developing embryos were staged in incubator at 28°C according to Kimmel *et al*. (29). Before any procedure, embryos were anesthetized in 0.02% tricaine.

### Cell culture, staining and microinjections

The HCT 116 human colorectal carcinoma cells were cultured in McCoy’s 5A Modified Medium supplemented with 10% fetal bovine serum (FBS), 10000 U/ml penicillin and 100 μg/ml streptomycin. The Mia Paca-2 human pancreatic carcinoma cells were cultured in DMEM supplemented with 10% FBS, 10000 U/ml penicillin and 100 μg/ml streptomycin. Cells were incubated at 37°C with 5% of CO_2_ in humidified atmosphere. Cells were detached at 80% confluence with 0.25% (w/v) trypsin-0.53 mM EDTA solution and stained with 10 μg/ml CM-Dil for 15 min at 37°C followed by 15 min on ice in darkness. Cells were washed and centrifuged three times by D-PBS and resuspended in D-PBS supplemented with 10% FBS to a final concentration of 100 cells/nl. All the reagents were supplemented by Thermo Fisher Scientific, Waltham, MA. Dechorionated embryos at 2 days post fertilization (dpf) were anesthetized and injected with four nanoliters of cells suspension in the left side of the perivitelline space using a heat-pulled needle and the PV830 Pneumatic PicoPump microinjector. The embryos were incubated at 35°C, and one hours after injection were screened with fluorescence microscope.

### Human tissue preparation and transplantation into zebrafish embryos

The clinical study was approved by the “Comitato Etico Regionale per la Sperimentazione Clinica della Toscana - sezione AREA VASTA NORD OVEST” (09/11/2017, prot n 70213). Human material from surgical resected specimens was obtained from the Azienda Ospedaliera Pisana (Pisa, Italy) after written informed consent of the patients and approval of local Ethical Committee. Tumor tissue screened by the histopathologist (from Histopathology unit, Cisanello facility) was washed three times with RPMI supplemented with 10000 U/ml penicillin, 100 μg/ml streptomycin and 100 μg/ml Amphotericin and cut into small pieces (1-3 mm) using a scalp blade. The pieces were then transferred to a 5 ml tube, and stained with either 40 μg/ml DiO in D-PBS (in case of esophageal and gastric cancers) or 40 μg/ml CM-Dil in D-PBS (in case of hepato-biliary-pancreatic cancers and intestinal cancers). The tissue pieces were incubated for 15 min at 37°C and 15 min in ice cube. Tissue pieces were then washed and centrifuged three times by D-PBS and resuspended in D-PBS supplemented with 10% FBS. For tissue transplantation we used the manual method proposed by Marques *et al.* 2009 (12). In particular before transplantation, small pieces of stained tissue were further disaggregated using Dumont forceps (No.5) into a relative size of 1/4 to 1/2 the size of the yolk. Tissue pieces with the correct size were transferred to 1% agarose disks in multiwell plates in which the 2 dpf embryos were laying, ready for transplantation. A glass transplantation needle was used to transfer the tissue into the yolk. The tissue was picked up, put on top of the yolk and then pushed inside. The yolk usually sealed itself and in the majority of embryos, the tumor remained in the yolk. After transplantation, embryos were incubated for 2 h at 35°C, then embryos were checked for presence of tissue and incubated at 35°C in E3 supplemented with 10000 U/ml penicillin and 100 μg/ml streptomycin with the presence or absence of drugs for the following days in the respect of the treatment plan.

### Anticancer drugs toxicity and treatment plan

Groups of 30 embryos (2 dpf) arrayed in multiwell plates were exposed to E3 supplemented with 10000 U/ml penicillin and 100 μg/ml streptomycin unmodified (control) and modified with the chemotherapy drug at 35°C for 24h added with increasing concentrations (Tables S1, S2).

The drugs were refreshed each day for the three days of treatment plan. 3 days after treatment (3 dpt) zebrafish larvae were fixed in 4% paraformaldehyde in PBS at 4°C over night. After that, they were dehydrated with increasing concentration of ethanol, and analyzed by stereo microscope to evaluate the phenotype (normal, death, aberrant).

### Microscopy and efficacy evaluation

Two hours post injection (2 hpi) zebrafish embryos xenotransplanted with cancer cell lines were anesthetized with 0.02% tricaine and positioned laterally, with the site of the implantation to the top. The embryos were imaged by fluorescence microscope and transferred to a 24-well plate (one embryo/well) containing chemotherapy compounds in E3 supplemented with 10000 U/ml penicillin and 100 μg/ml streptomycin or E3 supplemented with 10000 U/ml penicillin and 100 μg/ml streptomycin unmodified (control). All embryos were imaged everyday during the time course of the treatment. The size of the tumor area was measured by using ImageJ.

### Histopathology

At 2 dpi (4 dpf) the xenografted larvae were fixed in 4% paraformaldehyde for 1h at room temperature, followed by paraffin embedding for hematoxylin & eosin staining or OCT embedding for Hoechst staining. Larvae were respectively sectioned with microtome or cryostat, along the sagittal plane at a thickness of 8 μm.

Histopathological analysis was performed on paraffin sections stained by hematoxylin & eosin (Merck KGaA, Germany) and digitally imaged using Nikon Eclipse E600 microscope.

Cryostat sections were Hoechst 33342 counterstained. Digital images of the stained sections were generated using a Nikon Eclipse Ti. Pyknotic cells was counted at 40 X magnification within the epifluorescence DAPI image.

### Statistical analysis

We used GraphPad Prism 7 as statistical analysis software. Data analysis was performed by ANOVA, followed by Bonferroni correction or Dunnett’s post-hoc test or t-test. Statistical significance was set to 5%.

## ACKNOWLEDGMENT

The authors thank Ida Montesanti and Noemi Nardillo for supporting zebrafish imaging and Dr Martina Giannaccini for the experimental advices. This work was supported by Fondazione Pisa (project 114/16).

